# Negative autoregulation matches production and demand in synthetic transcriptional networks

**DOI:** 10.1101/000430

**Authors:** Elisa Franco, Giulia Giordano, Per-Ola Forsberg, Richard M. Murray

**Author notes:** University of California at Riverside (Elisa Franco). University of Udine (Giulia Giordano). California Institute of Technology (Richard M. Murray).

## Abstract

We propose a negative feedback architecture that regulates activity of artificial genes, or “genelets”, to meet their output downstream demand, achieving robustness with respect to uncertain open-loop output production rates. In particular, we consider the case where the outputs of two genelets interact to form a single assembled product. We show with analysis and experiments that negative autoregulation matches the production and demand of the outputs: the magnitude of the regulatory signal is proportional to the “error” between the circuit output concentration and its actual demand. This two-device system is experimentally implemented using *in vitro* transcriptional networks, where reactions are systematically designed by optimizing nucleic acid sequences with publicly available software packages. We build a predictive ordinary differential equation (ODE) model that captures the dynamics of the system, and can be used to numerically assess the scalability of this architecture to larger sets of interconnected genes. Finally, with numerical simulations we contrast our negative autoregulation scheme with a cross-activation architecture, which is less scalable and results in slower response times.

## 1 Introduction

Our increased understanding of biological parts enables their use in a variety of new applications^1^ of growing complexity, ranging from nanofabrication to drug production and delivery. When a large number of molecular devices are required to operate together within a system to achieve an overall functionality, it is essential that the output of each device is automatically tuned to meet its demand. For instance, poorly regulated production of an exogenous protein in a synthetic circuit may cause lethal host overloading;^2–4^ similarly, mismatched concentrations of RNA species forming a self-assembled structure, where simultaneous stoichiometric transcription of components is required, may result in the formation of undesired complexes and incorrect assemblies, both *in vitro*^5^ and *in vivo*.^6^ In other words, the functionality of a large scale synthetic system may deteriorate if the input/output behavior of individual synthetic genes or pathways characterized in isolation does not automatically meet specifications in its network context. Rather than fine tuning a device to fit a range of contingent network demands, it is desirable to identify design principles that would automatically ensure a demand-adaptive operation.

In traditional engineering fields, the challenge of adapting the output behavior of a device to reach the desired operating point is met by routinely employing negative feedback at a variety of scales (from individual transistors to layered network control systems). Consider, for instance, a device *S* whose output *y* is required to track a reference *r* (Figure 2A). A negative feedback loop causes the input to the regulated process *S* to be proportional with opposite sign to the error *e* between the output *y* and the (possibly changing) reference value *r*. Thus, the system’s response is always driven by an input with opposite trend relative to the error *e*. If, for instance, *y* exceeds *r* the error is positive, but the input to *S* is negative and drives “down” the response of *S*. In addition to maintaining a desired output level, negative feedback generally gives us the ability to redesign the dynamics of a system, and improve its robustness with respect to parametric uncertainty.^7^

**Figure 1:**
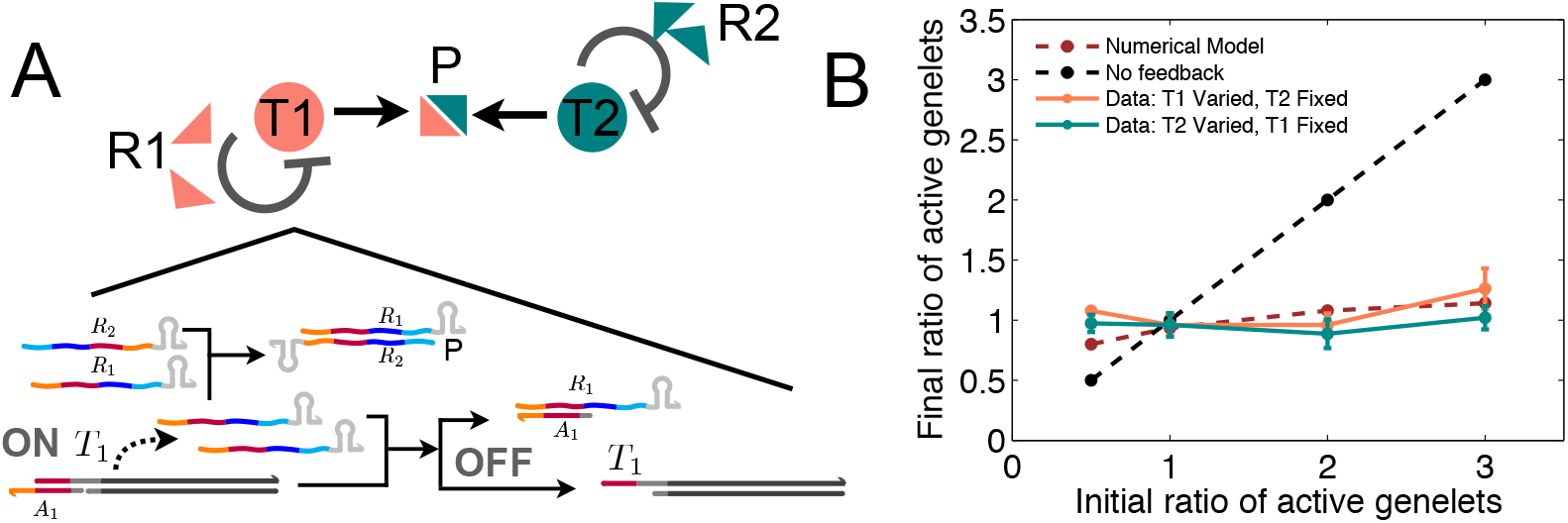
A: Schematic of our negative feedback scheme and its implementation using synthetic transcriptional “genelets”. B: Experimental data showing that activity levels of the genelets are matched, achieving balanced production and demand of the RNA species forming product P.

**Figure 2:**
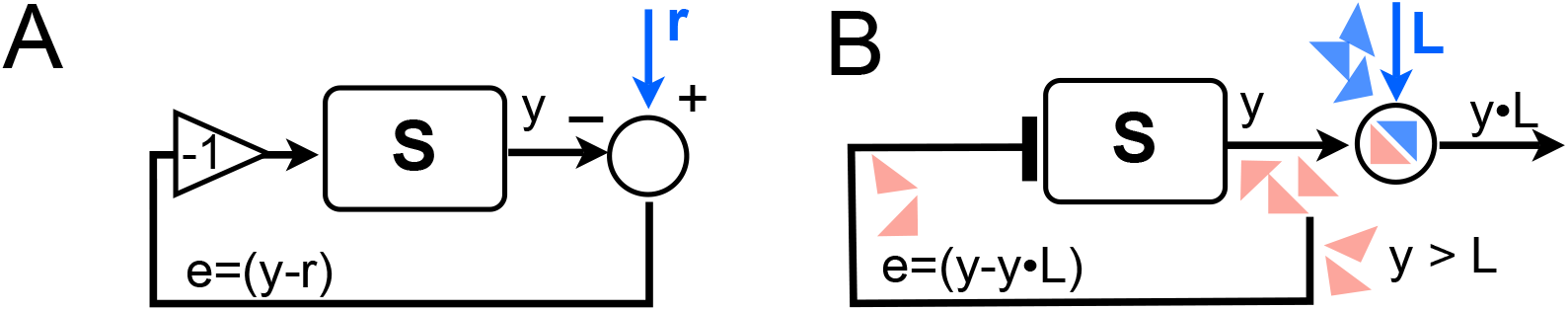
A: General structure of a negative feedback loop, where the system input counteracts the error between the desired and actual system output. B: Negative feedback scheme for a molecular system, where an excess production of y is used to downregulate the “activity” of the system.

Negative feedback is ubiquitous in biomolecular networks. For example, negative autoregulation is a motif present in over 40% of genes in *E. coli*.^8, 9^ This mechanism is associated with proteins that are generally in low demand,^10, 11^ and reduces noise^12–14^ and mutation rates^15^ in gene expression profiles. In the context of synthetic biology, negative autoregulation has been used to achieve faster response speed^16^and to improve robustness.^12, 15^ The development of novel, tunable repression mechanisms promises to improve our ability to control dynamics and manage noise of increasingly complex molecular circuits both in cellular hosts^17, 18^ and in cell-free systems.^19–21^ However, the use of negative feedback to match production and demand within a biochemical reaction network has, to our knowledge, not been demonstrated.

In this paper we propose to use negative feedback to accurately regulate activity of components so they can meet their output downstream demand, achieving robustness with respect to uncertain open-loop (*i. e.* in the absence of feedback) output production rates. Figure 2B shows a scheme of this feedback architecture, which closely mimics the structure of a typical negative feedback circuit in electrical or mechanical systems. The output *y* of component *S* binds to a downstream target *L*, which represents the demand for *y*; we design a negative feedback pathway to use excess *y* (not bound to *L*) to reduce its own production rate: thus, the magnitude of the regulatory signal is proportional to the “error” between the circuit output concentration and its actual demand. If in turn *L* is the output of another circuit, it is conceivable that a negative feedback loop in each individual circuit would help matching production and demand in the overall system. With analysis and experiments we show that negative autoregulation yields matching output fluxes for both circuits.

The two-circuit system is implemented using *in vitro* transcriptional networks,^19, 22, 23^ a versatile toolbox to program and implement dynamic behaviors in nucleic acid reaction networks. Within the general context of cell-free systems,^24^ this platform allows to rapidly engineer molecular functions in a controlled environment with reduced uncertainty. We designed two synthetic genes to transcribe RNA outputs that bind to form a complex; each RNA species is also designed to downregulate its own production through promoter displacement.^19^ Thus, excess of either species modulates the genes’ activity and achieves matched promoter activity levels. The product formation reaction and the inhibitory pathways are systematically engineered by optimizing nucleic acid sequence complementarity domains, using publicly available software packages.^25, 26^ We build a predictive ordinary differential equation (ODE) model that captures the dynamics of the system, and can be used to numerically assess the scalability of this architecture to larger sets of interconnected genes. Finally, with numerical simulations we contrast the performance of our negative autoregulation scheme with the behavior of a cross-activation architecture, which is less scalable and results in slower response times. This work builds on preliminary numerical analysis and experiments on transcription matching synthetic systems.^27–30^ We foresee that systematic use of similar negative feedback architectures will play a major role in the scalability of *in vitro* biomolecular systems, including logic,^31^ dynamic,^23^ and self-assembly networks.^5, 32^

## 2. Results

### 2.1 Negative feedback can modulate activity to meet downstream demand

We begin by considering a simple model problem: a molecule *R* is produced by species *T*, and binds to a target *L*:

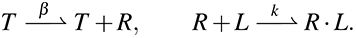

These reactions may represent, for instance, RNA or protein production followed by binding of the product to a downstream binding site or ligand. In the absence of any regulatory pathway feeding back to *T* information regarding the effective “consumption” of *R* by the target *L*, the production and demand of *R* are not automatically matched: thus, an excess of unused *R* may accumulate in solution for regimes where the demand does not exceed maximum production rates. However, if we program a reaction

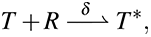

whereby species *T* bound to *R* becomes an inactive species *T*\*, we introduce a negative feedback mechanism that is proportional to the unused amount of [*R*] ∝ [*R^tot^*] − [*R · L*], thus proportional to the error between production and demand. The scheme is represented in Figure 3 A. Assuming that the concentration of the “demand” species *L* is constant, that the total amount of *T* is constant ([*T*] + [*T*\*] = [*T^tot^*]), and finally that inactive *T*\* spontaneously reverts to its active state at a certain rate 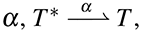 the system is described by the following set of ODEs:

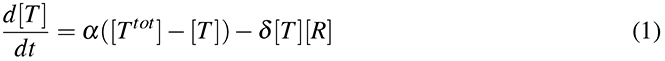

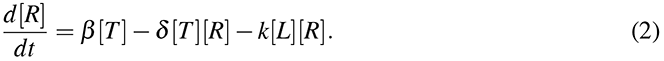

**Figure 3:**
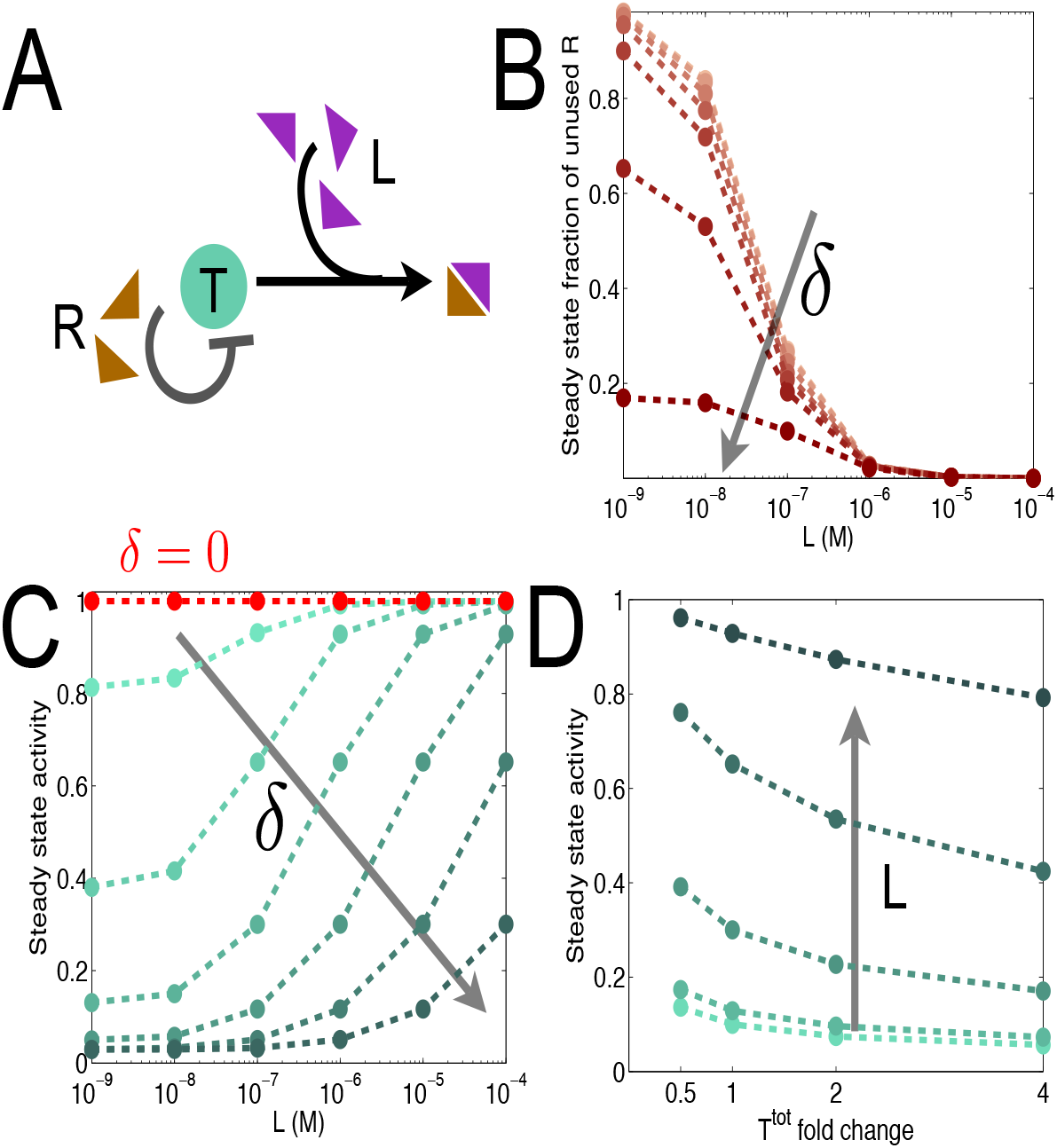
A: Scheme of negative feedback where output *R* not bound to the target load *L* is used to downregulate its production. B: The steady state fraction ([*R^free^*]/[*R^tot^*]) of unused *R* as a function of the downstream load is reduced using a high negative feedback rate *δ*. In Panels B and D, we considered *δ* = 0, 5, 50, 5 · 10^2^, 5 · 10^3^, 5 · 10^4^, 5 · 10^5^/M/s. In Panel D, the nominal concentration of [*T^tot^*] is 100 nM, and the load was varied as [*L*] = 1, 10, 10^2^, 10^3^, 10^4^ nM.

For illustrative purposes we numerically simulate these differential equations, choosing nominal parameters [*T^tot^*] = 100 nM, *α* = 3 · 10^−4^/s, *β* = 0.1/s, *k* = 2 · 10^−3^/M/s, *δ* = 5 · 10^2^/M/s; these concentrations and rates are within a realistic range for *in vitro* reaction systems.^23, 24^ In Figure 3 we explore the steady state behavior of the system as a function of the feedback parameter *δ*, the total amount of load *L*, and the total concentration of generating species *T*. First, as shown in Figure 3B, we note that a suitably high feedback rate *δ* reduces the steady state fraction of unused output [*R*]/[*R^tot^*] (output not bound to its load): this means waste in the system is reduced. In addition, for a given, large *δ*, a significant variation in load results in a moderate variation in the fraction of unused output: this behavior is consistent with the role of high feedback in reducing load sensitivity in retroactivity theory.^33^ In Figure 3C we observe that in the presence of feedback the activity of the generating species, defined as [*T*]/[*T^tot^*], is modulated by the demand *L*. Finally, Figure 3D shows that the presence of negative feedback yields closed loop activity levels that (given a certain demand) are robust with respect to uncertainty in [*T^tot^*], which is a simple open loop knob to scale the production rate of *R*.

### 2.2 Matching output fluxes in interconnected devices

In many practical cases, several molecular species in a network bind stoichiometrically to form an overall product. For instance, these species could be RNA strands^5^ or proteins^34^ self-assembling in a nanostructure. To avoid excess production and accumulation of any participating species we can use the negative feedback scheme described above. For simplicity, we begin by considering a network where two generating species *T*_1_ and *T*_2_ produce assembling outputs that self-inhibit according to the following reactions:

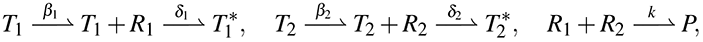

where *P* is an assembled product, and again we assume that the total amount of the generating species is conserved, [*T*_*i*_^*tot*^] = *T_i_* + *T*_*i*_^*^, *i* = 1, 2. The dynamics of [*T*_1_] and [*T*_2_] are thus described by ODEs identical to equation (1), while the dynamics of [*R_i_*] become:

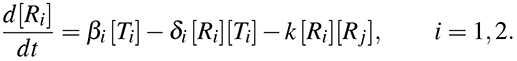

Example solutions to these ODEs are shown in Figure 5A, where we chose identical parameters for the two subsystems consistent with our previous simulations at Figure 3 (*α*_1_ = *α*_2_ = *α* = 3 · 10^−4^/s, and similarly defined *β* = 0.1/s, *k* = 2 · 10^−3^/M/s, *δ* = 5 · 10^2^/M/s).

**Figure 4:**
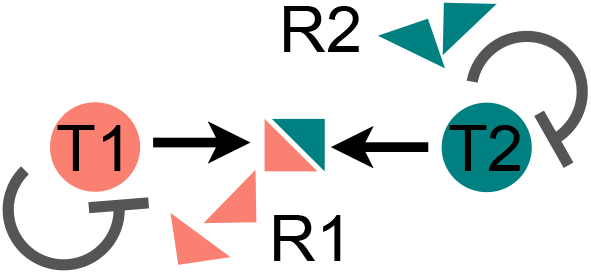
Our two-device negative feedback architecture.

**Figure 5:**
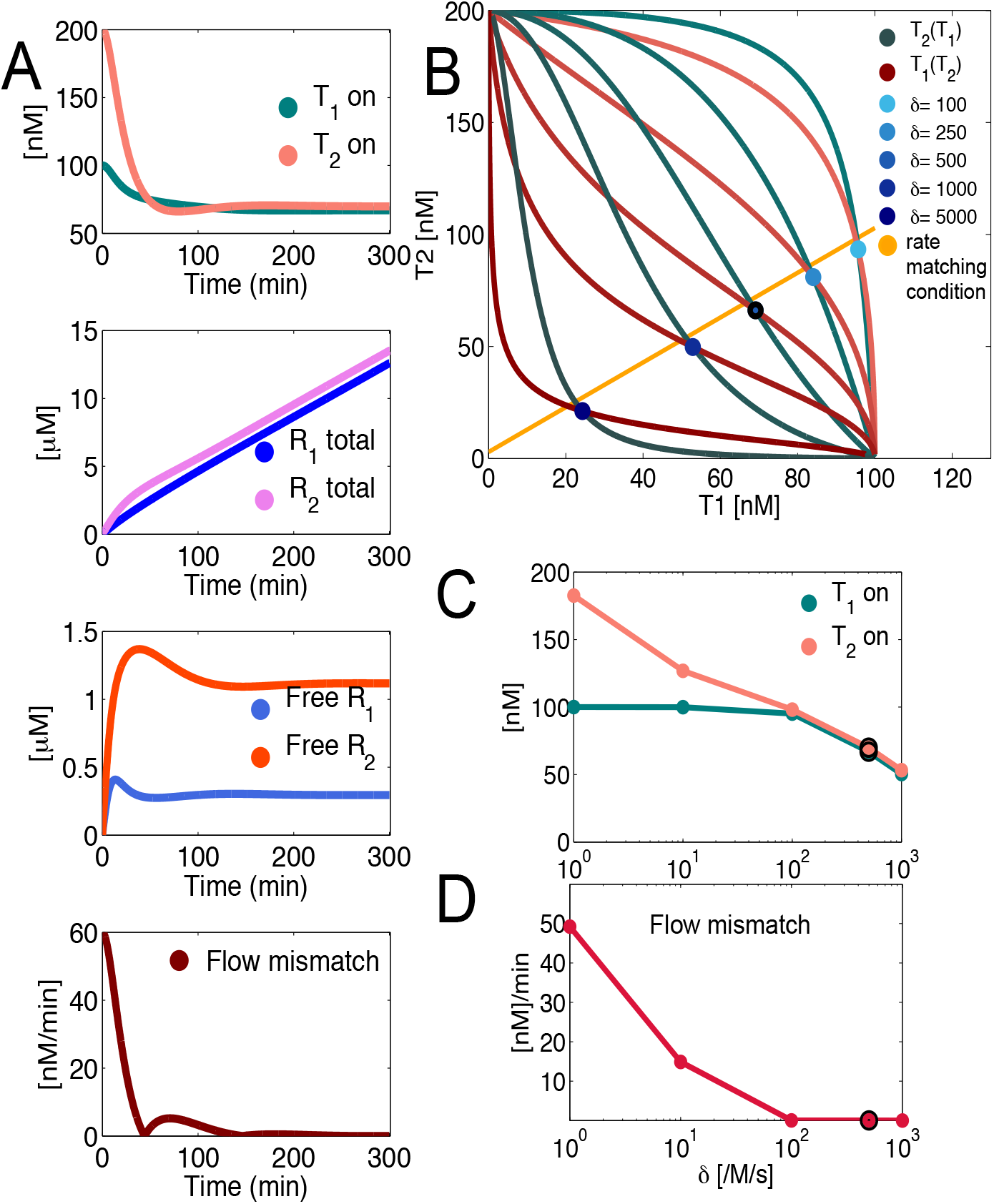
A: Numerical simulations showing example trajectories for the two-component negative feedback architecture. From top to bottom: time course of active *T*_1_ and *T*_2_; time course of total produced outputs *R*_1_ and *R*_2_; time course of unbound *R*_1_ and *R*_2_; time course for the flux mismatch in the production of total *R*_1_ and *R*_2_. B: Nullclines for *T*_1_ and *T*_2_ for a range of values of *δ*, and flux matching condition in equation (3) (orange). C: Steady state activity of *T*_1_ and *T*_2_ as a function of the negative feedback parameter *δ*. D: Mismatch in the flux of *R*_1_ and *R*_2_ as a function of *δ*. The dark circle in panels C and D marks the nominal conditions used for the time courses plotted in panel A.

Expressions for the nullclines of the system are derived in Section 2.1 of the Supplementary Information (SI), and the equilibria (intersections of the nullclines) are numerically evaluated as a function of the negative feedback reaction as shown in Figure 5B. At steady state, the concentration of active *T*_1_ is nearly identical to the active concentration of *T*_2_. Figure 5C, however, shows that this property breaks down when the negative feedback rate *δ* is too low; while a high *δ* guarantees matched activity levels, it also causes an overall lower activity level for the system and pushes down the production of *P* = *R*_1_ *· R*_2_ complex.

We also ask if, at a stationary regime, the dynamic behaviors of *R*_1_ and *R*_2_ is similar. We find that the flux of both outputs are identical when the active concentrations [*T̅*_1_] and [*T̅*_2_] are related as follows (*cf.* Section 2.1 of the SI):

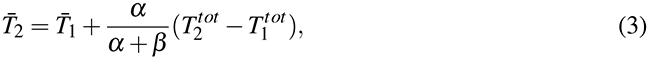

where for simplicity we assumed *α*_1_ = *α*_2_ = *α*, *β*_1_ = *β*_2_ = *β*, and *δ*_1_ = *δ*_2_ = *δ*. Thus, when *β* is sufficiently large relative to *α*, the flux of the two outputs is matched (orange line in Figure 5B). Numerically, we observe again that flux matching is lost for low values of *δ* (Figure 5D).

#### 2.2.1 Experimental results: Negative autoregulation balances RNA transcription rates in a two-gene artificial network

We implemented experimentally the two-species model problem described above using *in vitro* transcriptional circuits.^19^ A sketch of the reactions for subsystem 1 is in Figure 6, where we highlight the regulatory domains of nucleic acid species, the main chemical reactions occurring, and the simple model pathways they correspond to. Two short, linear genetic switches, or genelets, correspond to species *T*_1_ and *T*_2_, whose RNA transcripts are the outputs *R*_1_ and *R*_2_. Transcription is carried out by T7 RNA polymerase. The transcripts are designed to bind and form an inert RNA complex *P*. (Since the focus of this work is the investigation of the effects of feedback, the structure of *P* and its functionality as a stand alone complex are neglected.) Genelets have a nicked T7 bacteriophage promoter sequence which can be displaced by toehold-mediated branch migration.^35^ We design the RNA output of each genelet to be complementary to the portion of the promoter that can be displaced (activator strand *A_i_*): therefore, free RNA in solution displaces the activator and self-inhibits its own production bringing the genelet in an “off” state. Degradation in the system is introduced by RNase H, which hydrolizes RNA in DNA/RNA complexes. DNA strands were systematically designed by thermodynamic analysis using the Winfree lab DNA design toolbox for MATLAB, Nupack,^26^ and Mfold.^36^ Sequences were optimized to yield free energy gains favoring the desired reactions, and to avoid unwanted secondary structures and crosstalk. For example, we ensured that the *R*_1_*R*_2_ complex formation reaction be more favorable than the self-inhibition reaction: because roughly twice as many base-pairs are complementary in the *R*_1_*R*_2_ complex relative to the *R_i_A_i_* (inhibition) complex, the Δ*G* of formation of *R*_1_*R*_2_ is *≈* −110 kcal/mol, twice as large (in absolute value) as the Δ*G* of formation of *R_i_A_i_*, which is *≈* −41 kcal/mol. Strand sequences and complete reaction schematics are in Section 1 of the SI.

**Figure 6:**
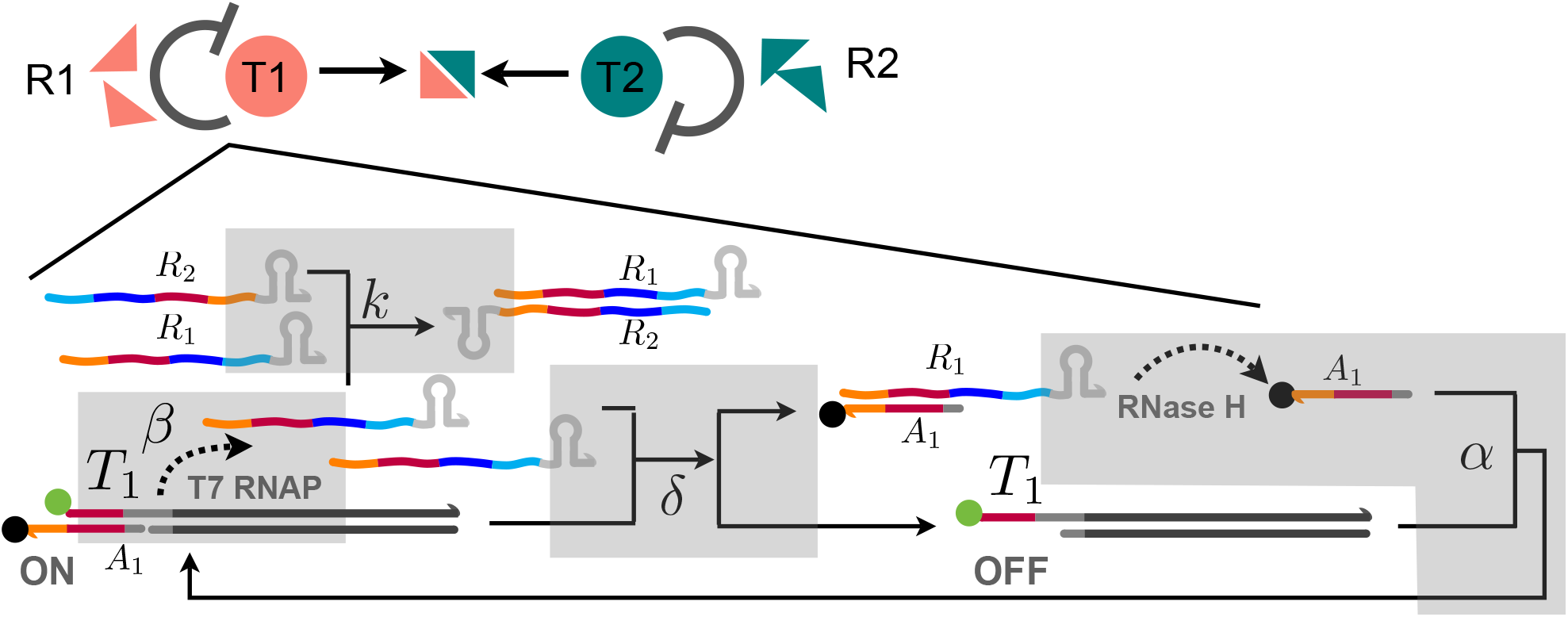
Summary scheme of DNA species and enzymes used to implement experimentally our negative feedback system for RNA production matching. Only subsystem 1 is represented (subsystem 2 is specular to subsystem 1). Complementary domains are indicated with the same color. RNA species *R*_1_ and *R*_2_, transcribed by active genelets *T*_1_ and *T*_2_, are designed to be complementary (dark red and dark blue domains), but also to function as self-inhibiting species. The orange-dark red domains in *R*_1_ indicate complementarity to the nicked portion of the promoter, activator *A*_1_, which is displaced by free *R*_1_ (in excess with respect to *R*_2_) through toehold-mediated branch migration. The complex *R*_1_*A*_1_ is degraded by RNAse H, which releases in solution *A*_1_; thus, *A*_1_ and *T*_1_ bind, recovering the genelet activity. Genelet activity can be tracked using a fluorophore-quencher pair (green and black dot positioned on *T*_1_ and *A*_1_). Gray boxes map the main pathways in this system to the simplified reactions of our model problem.

We expect the feedback scheme to downregulate the production of either RNA species when in excess with respect to the other. For instance, if the concentration of genelet 1 is twice the concentration of genelet 2, in the absence of regulation the concentration of *R*_1_ produced will clearly exceed that of *R*_2_. However, in the presence of negative feedback, we expect to observe downregulation of the active gene 1 to achieve concentrations close to the active concentration of gene 2. This expectation is quantitatively plausible, since the promoters used in both genelets are identical and their activity is thus similar. We can easily verify this hypothesis by labeling the 5′ end of the non-template strand of each genelet with a fluorescent dye, and by labeling the corresponding activator strand with a quencher on the 3′ end. Inactive templates will emit a high fluorescence signal, while the signal of active templates will be quenched (Figure 6, green and black dots respectively represent fluorophores and quenchers). For instance, when *A*_1_ is stripped off active *T*_1_, the *T*_1_ fluorescence signal will increase. However, fluorescence traces reported here are processed to show a high measured signal in correspondence to a high genelet activity. In our experiments the total amount of activators is stoichiometric to the total amount of templates; for brevity we will just indicate the total concentration of *T^tot^*, with the understanding that [*A*_*i*_^*tot*^] = [*T*_*i*_^*tot*^].

Figure 7A shows the behavior of the system in the scenario described above, i.e. when the total concentration of the two genelets is in a 2:1 ratio. As soon as enzymes are added in solution and transcription is initiated, the formation of complex *R*_1_*R*_2_ is limited by the lower production rate of gene 2 (present in a lower amount). Thus, excess *R*_1_ reduces its own production by displacing its activator from the genelet, and balances the active concentration of the two genes to be practically identical. Thus, the steady state ratio of active genelets is close to one. Dashed lines in the figure are numerical traces generated by a detailed model comprised of several differential equations, whose parameters were fitted to the collected data.

**Figure 7:**
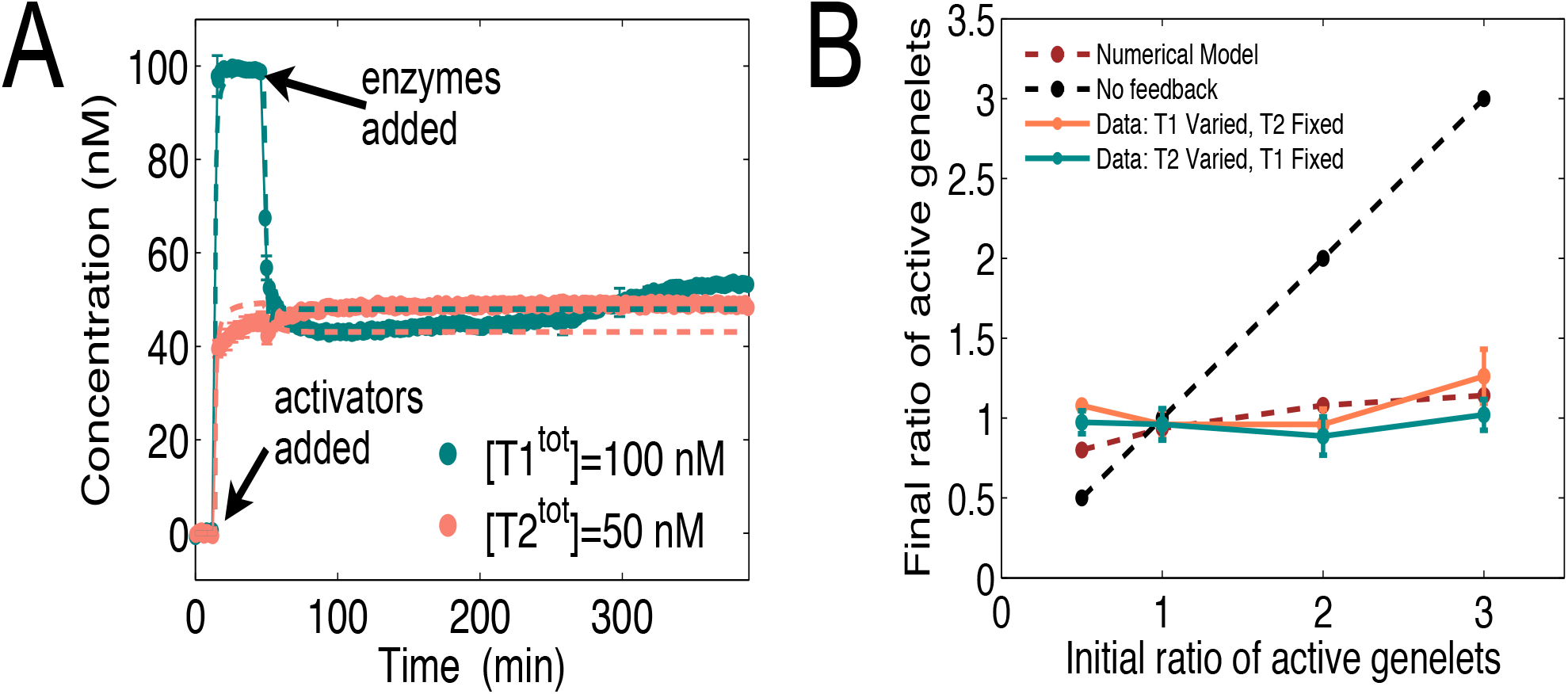
A: Experimental fluorimetry data showing a typical time course of our system. Experiments were run in triplicates. Once activators are added, both genelets become fully active. Addition of enzymes initiates production of *R*_1_ and *R*_2_, which rapidly form a complex; excess of either species is expected to downregulate its own genelet activity. In this case, [*T*_1_^*tot*^] is present in solution at a concentration which is twice that of [*T*_2_^*tot*^]: as expected, excess *R*_1_ inactivates *T*_1_ to activity levels comparable to [*T*_2_^*tot*^]. B: We screened the outcome of various time courses (panel A), where we varied the total concentration of genelets and measured the steady state activity of each genelet. This plot summarizes our results, showing that in a wide range of conditions the steady state activity of the genelets always achieves a 1:1 ratio, thus matching production and demand of the RNA outputs.

We repeated this experiment for a variety of genelet ratios, keeping the concentration of one of the genelets constant and varying the concentration of the other gene. The steady state ratio of the active genelets was close to one in all cases (our complete data sets are in Section 1.6 of the SI). Figure 7B summarizes this experimental assay and shows that our negative autoregulation scheme guarantees matched production and demand in a wide range of conditions.

When the concentration of genelets varies over time, the negative feedback scheme handles a change in demand by automatically adapting the amount of each active genelet. Figure 8A shows that abrupt changes in the total concentration of one of the genelets are followed by an adjustment in the concentration of the excess species to guarantee a matched flux of the RNA products. We estimated the total amount of each RNA species produced during this experiment by gel electrophoresis, verifying that their production rate is adapted and their concentration is in a 1:1 stoichiometry.

**Figure 8:**
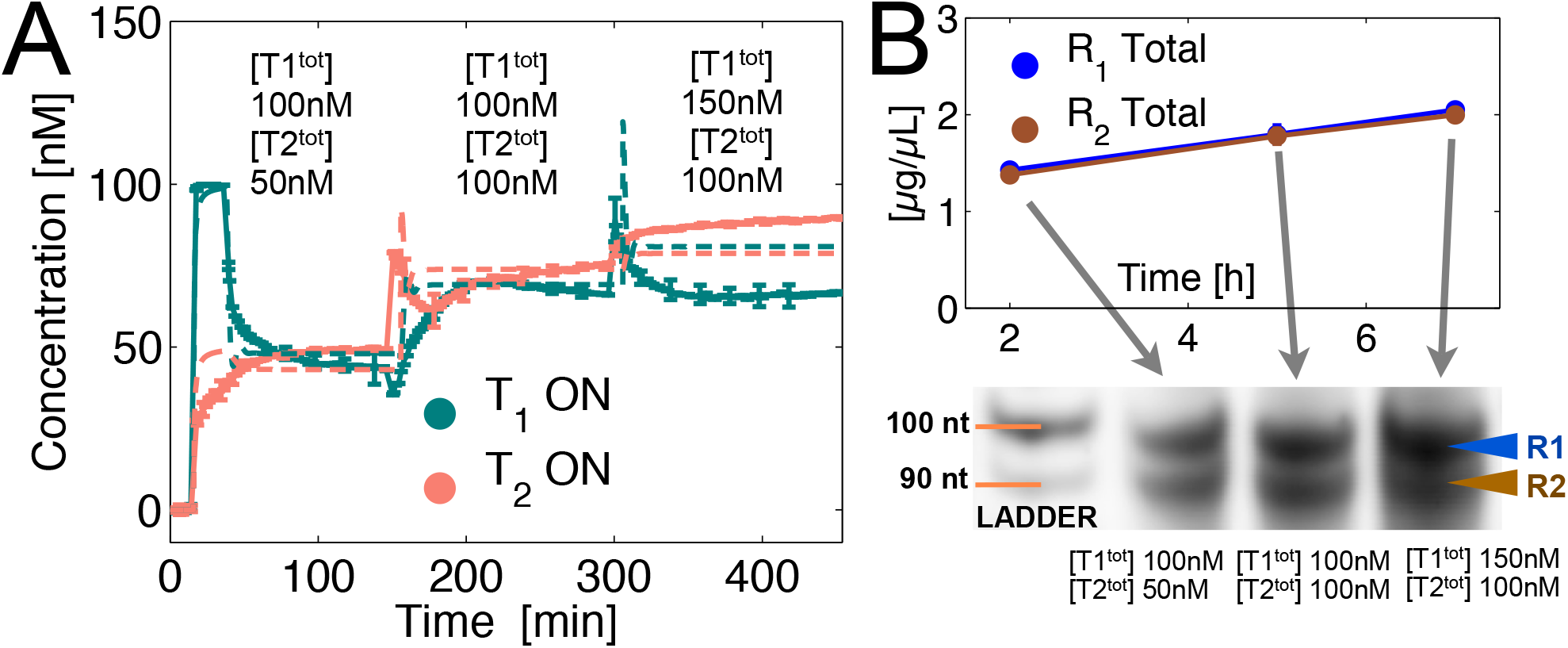
A: We varied the total concentration of genelets over time, maintaining activators and templates stoichiometric. Experiments were run in triplicates. The system shows adaptation: when the concentration [*T*_2_^*tot*^] is increased to 100 nM, we observe an increase in the activity level for *T*_1_, which was previously half-repressed. Further increase in the concentration [*T*_1_^*tot*^], however, only marginally changes the activity levels, because the activity of *T*_2_ is nearly at maximal levels. B: We sampled our time course experiments over time, and estimated the concentration of *R*_1_ and *R*_2_ through gel electrophoresis. The two concentrations remain comparable despite the changes in total genelet concentrations, further supporting our hypothesis that this negative feedback scheme matches production and demand by regulating genelet activity.

#### 2.2.2 Mathematical modeling

We built a model for the *in vitro* two-gene flux matching system, starting from a complete list of reactions involving the nucleic acid and enzyme species. Using the law of mass action, we derived a set of ordinary differential equations (ODEs) which were numerically solved using MATLAB. The list of reactions (reported in Section 2.2 of the SI) includes both the designed interactions among species, and some of the expected undesired reactions. Specifically, we include reactions of (weak) transcription for genelets in an off state. In addition, our design specifications result in an undesired binding domain between T_i_ and R_j_, which is considered a further off state of the genelet. Such complex is a substrate for RNase H and the RNA strand is degraded by the enzyme, releasing the genelet activation domain. The transcription efficiency of an RNA-DNA promoter complex is very low.^23^ We are aware of other sources of uncertainty when modeling genelet systems, including transcription bursting and RNA polymerase activity decay phenomena, abortive transcription, and partial RNase H mediated degradation of RNA-DNA hybrids (resulting in the accumulation of short RNA species). We found that these events play an important role in complex dynamical systems such as oscillators,^22, 23^ whose temporal behavior is highly sensitive to variations in the enzyme characteristics (which change from batch to batch) and notoriously difficult to model quantitatively. However, the experimental outcomes of our negative autoregulation system were satisfactorily captured by a detailed model that did not include the aforementioned phenomena.

### 2.3 Scalability and alternative architectures

The size of synthetic biological circuits, from metabolic networks^37^ to molecular computers,^31, 38^ is rapidly increasing to include hundreds of components. Thus, we ask if our negative feeback scheme is scalable to a larger number of interconnected components. For instance, our two-gene circuit, where two RNA outputs interact to form a complex, could be extended to *n* genes whose outputs assemble in a single product. From a practical perspective, formation of co-transcriptional self-assembled RNA structures have been demonstrated^5^ in the absence of any regulatory pathways for transcription; the introduction of feedback could improve the stoichiometry of RNA components, and thus improve the yield of correctly assembled structures.

We also ask if alternative feedback mechanisms can achieve production and demand matching in molecular devices. Positive feedback can easily generate instability in conventional engineered systems, and is thus carefully avoided by systems and control engineers. In contrast, positive feedback is commonly found in biology, in particular in gene networks^9^ in the context of autoregulation^10^ or within more complex motifs.^39, 40^ Motivated by Savageau’s theory of positive autoregulation being common for proteins in high demand,^10, 11^ we consider an alternative architecture for matching production and demand that is based on cross activation.

#### 2.3.1 Scalability

To investigate how the performance of negative autoregulation would scale in the context of a network composed by *n* molecular devices, we identified three canonical topologies for the output interactions. We say that two devices are interconnected if their outputs bind or assemble to form one or more products. Our two-gene network can immediately be scaled up to what we can call a “single product” topology (Figure 9A) with *n* participating species. When more than one assembled products is generated, we identify two limit cases: the output of each device participates in at most two products, creating a “neighbor” topology (Figure 9B); the output of each device participates in *n* − 1 products, generating a “handshake” topology (Figure 9C). From an input/output perspective, we expect that the neighbor and handshake topologies can be rendered equivalent to the single product architecture, by designing appropriate downstream interactions among the network complexes. For example, complexes created by interacting pairs of outputs (neighbor topology) may further interact with one another and generate a single output assembly (single product).

**Figure 9:**
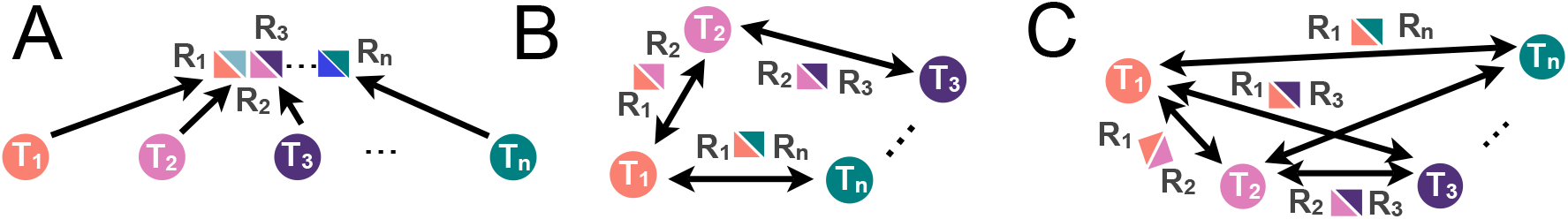
We explore the scalability of our two-device network by looking at three limit cases where *n* devices are interconnected through their binding outputs. A: Single product interconnection. B: Neighbor interconnection. C: Handshake interconnection.

We ask if, in all these possible topologies, our negative autoregulation scheme can still help modulating the activity of each device in order to match production and demand of each output. With numerical simulations we explored the behavior of up to four-component networks for each of the topologies described above. The simple model of ODEs (1)-(2) can be straightforwardly modified to model each topology, as reported in Section 3.1 of the SI.

Simulation results show that negative feedback is still effective in regulating the devices activity: it reduces both steady state activity of *T_i_* and the mean flow mismatch. The evolution over time of each species is very similar to the one shown in Figure 5A for the case of two molecular devices; example time trajectories for *n* = 4 are reported Section 3.1 of the SI.

From a network design perspective, it is interesting to explore the performance of different interconnection topologies as a function of key parameters such as the feedback strength, *δ*, and the rate of spontaneous gene activation, *α*. For illustrative purposes, in Figure 10 we compare the performance of our three feedback topologies for *n* = 4 within a range of values for *δ* and *α*. In each panel, a pink square marks the system behavior in nominal conditions, *k_ij_* = 2 · 10^3^/M/s for the handshake/neighbor topology, *k* = 6 · 10^3^/M/s for the single product topology, *δ_i_* = 5 · 10^3^ /M/s, *α_i_* = 3 · 10*^−^*^4^ /s, *β_i_* = 1 · 10*^−^*^2^ /s. An imbalance in the production rates of *R_i_* is created by setting [*T_i_*](0) = [*T*_*i*_^*tot*^], while [*R_i_*](0) = 0, choosing [*T*_1_^*tot*^] = 100 nM, [*T*_2_^*tot*^] = 200 nM, [*T*_3_^*tot*^] = 300 nM, [*T*_4_^*tot*^] = 150 nM. We report the percent steady state activity level of *T_i_*, defined as [*T_i_*]/[*T*_*i*_^*tot*^] · 100, and the flux mismatch for each pair of outputs: each point in these graphs corresponds to the behavior of each subsystem averaged over the last hour (stationary behavior) of a 10 hour numerical simulation. We also report the response time of *T_i_*, computed as the time it takes for the active *T_i_* trajectory to go from [*T_i_*(0)] − 10%Δ to [*T_i_*(0)] − 90%Δ, where Δ is the difference between its initial value [*T_i_*(0)] and its steady state value.

**Figure 10:**
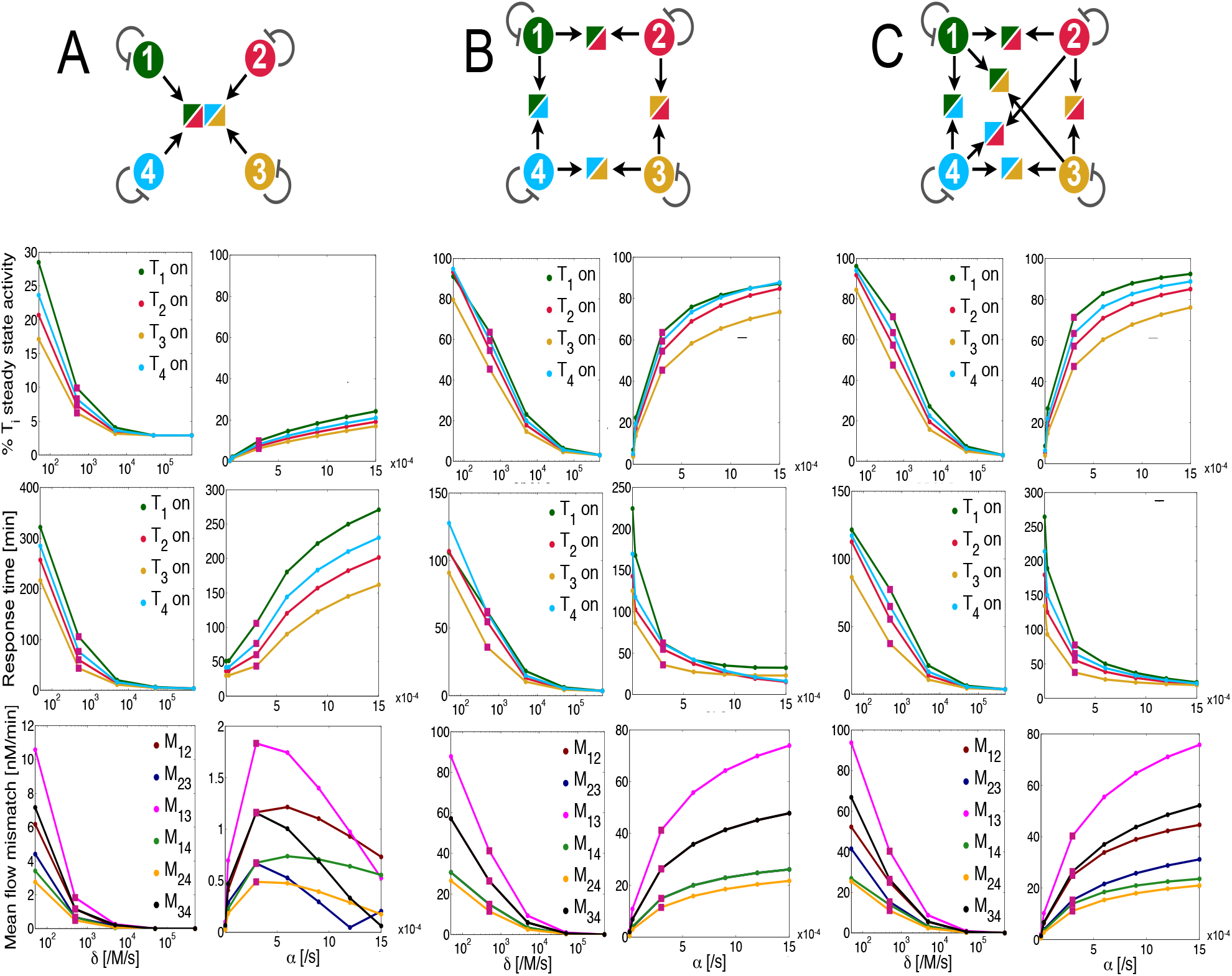
Sensitivity of *T_i_* percent activity, *T_i_* response time, and flux mismatch between pairs of outputs *R_i_*, with respect to the negative feedback rate *δ* and the spontaneous reactivation *α*. A: Single product interconnection. B: Neighbor interconnection. C: Handshake interconnection. Pink squares mark the system behavior in nominal conditions.

Referring to Figure 10, we can see that the steady state activity of *T_i_* is higher for neighbor (Figure 10 B) and handshake (Figure 10 C) topologies; nevertheless, for all topologies the steady state activity of *T_i_* decreases when *δ* increases, and it increases when *α* increases. The sensitivity of steady state *T_i_* with respect to *α* is lowest in the single product topology: this may be regarded as a benefit or a flaw of the system, depending on the overall product complex (*P* = ∏*_i_ R_i_*) downstream demand. The flux mismatch is most significantly reduced as a function of *δ* in the single product topology (Figure 10 A). However, this topology yields a much slower response time for *T_i_*, relative to the neighbor and handshake structures. The response time generally decreases in all topologies with large spontaneous gene activation rate *α*. However, the response time increases in the single product topology. Thus, while the single product topology is more effective in matching production and demand of each output *R_i_*, its response time is large relative to other topologies, and more sensitive to *α*.

#### 2.3.2 An alternative positive feedback architecture

We explore numerically the performance of a two-device system where excess outputs cross-activate their production, rather than self-inhibit. This scheme is expected to increase the overall network output production rate, due to mutual activation of the generating species. Figure 11 A shows a sketch of the system we consider. Two generating species *T*_1_ and *T*_2_ create outputs *R*_1_ and *R*_2_, which bind to form a product *P* = *R*_1_ *· R*_2_. Free molecules of *R_i_*, not incorporated in *P*, generate a positive loop by binding to inactive *T_j_* and activating it:

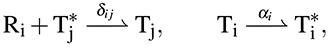

where again *T*_*i*_^*^ is an inactive complex and [*T*_*i*_^*tot*^] = [*T_i_*] + *T*_*i*_^*^. The total amount of *R_i_* is [*R*_*i*_^*tot*^] = [*R_i_*] + [*T_j_*] + [*P*]. We now assume that *T_i_* naturally reverts to its inactive state with rate *α_i_*. The corresponding differential equations are:

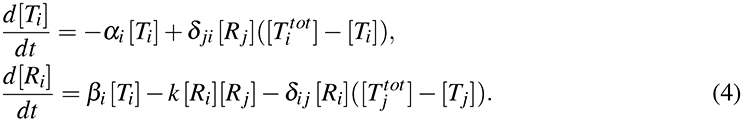

**Figure 11:**
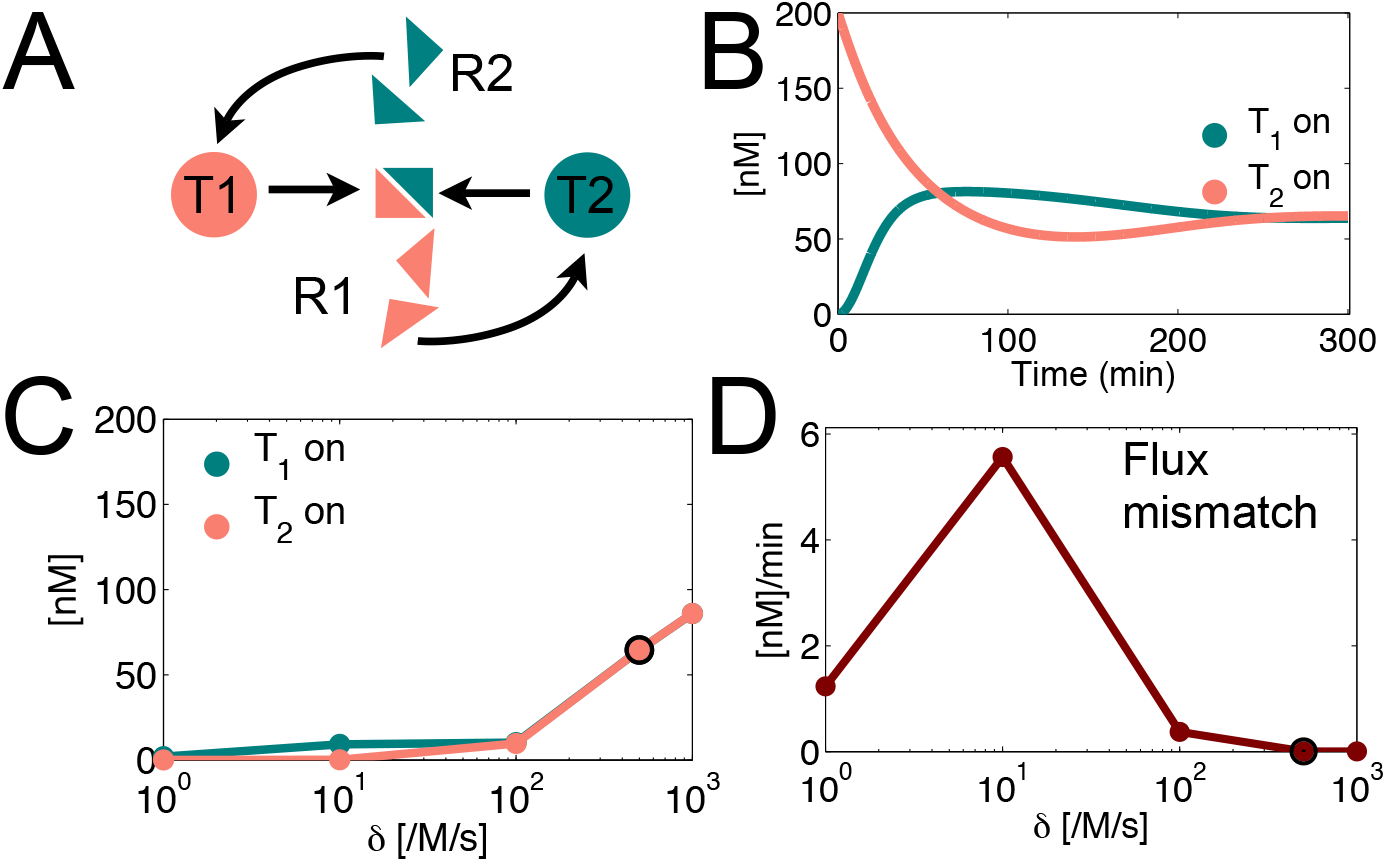
A: Positive feedback architecture to match production and demand of interconnected devices. B: Numerical simulation showing the time course of *T*_1_ and *T*_2_. C: Steady state activity of *T*_1_ and *T*_2_ as a function of the positive feedback parameter *δ*. D: Mismatch in the flux of *R*_1_ and *R*_2_ as a function of *δ*. The dark circle in panels C and D marks the nominal conditions used for the time course plotted in panel B.

The above differential equations were solved numerically. For illustrative purposes, our choice of parameters is consistent with the numerical study of the negative feedback circuit: *α*_1_ = *α*_2_ = 3 · 10*^−^*^4^ /s, *β*_1_ = *β*_2_ = 0.01 /s, *δ*_1_ = *δ*_2_ = 5 · 10^2^ /M/s, and *k* = 2 · 10^3^/M/s. The total amount of templates was chosen as [*T*_1_^*tot*^] = 100 nM, [*T*_2_^*tot*^] = 200 nM. The initial conditions of active [*T_i_*] are set as [*T*_1_](0) = 0 nM and [*T*_2_](0) = 200 nM, while [*R*_1_](0) = [*R*_2_](0) = 0. As a function of the positive feedback strength (for simplicity we picked *δ* = *δ*_1_ = *δ*_2_), the steady state amount of active *T_i_* clearly increases (we define our steady state as the mean active [*T_i_*] during the last hour of a 10 hours trajectory simulation), as shown in Figure 11 C. We compute flux of *R_i_* again as the derivative of the total amount of [*R*_*i*_^*tot*^] = [*R*_*i*_] + [*R*_*i*_*T_j_*] + [*P*]. The flux mismatch between *R*_1_ and *R*_2_ is defined again as the absolute value of the difference between the two fluxes; the average flux mismatch over the last hour of a 10–hour simulation is plotted as a function of *δ* in Figure 11 D. Unlike the negative feedback architecture (*cf.* Figure 5 D), the flux mismatch is not monotonically decreasing as a function of *δ*; however, a sufficiently large positive feedback yields matching fluxes and, as expected, higher activity levels relative to the negative feedback scheme (Figure 11 C and D).

We examined the nullclines and derived flux matching conditions for the positive feedback architecture as done for the negative feedback scheme; the complete derivations are in Section 4.1 of the SI. Again, we find that the circuit has, for a certain range of parameters, the ability to match the flux of outputs *R_i_* by upregulating the production of output in lack. Because the production rate of *R_i_* is limited by the finite maximal amount of activatable *T_i_* (whose maximal active concentration equals [*T*_*i*_^*tot*^]), the positive feedback loops cannot yield instability (*i. e.* uncontrolled increase) in the amount of unbound *R_i_*. However, we observed that an overall upregulation of *T_i_* activity results in slower response time for the circuit.

We explored the performance of the cross-activation scheme in the context of the larger-scale interconnection schemes considered in the previous section (Figure 9). First, we have to remark that a cross-activation scheme scales poorly with the number of devices in the network. The number of required regulatory reactions *n_reg_* is equal to the product of three factors: the number of devices *n*, the number *n_P_* of complexes generated by each device, and the number *n_r_* of reactions required to form each product (*n_reg_* = *nn_P_n_r_*). Thus, *n*(*n* − 1) regulatory reactions are required in the single product and handshake topologies, while 2*n* reactions are required in the neighbor topology. In contrast, the negative autoregulation scheme requires *n* regulatory reactions regardless of the chosen output interconnection topology. Nevertheless, we evaluated the performance of this scheme for a 3–devices network, for which handshake and neighbor topologies coincide. In Section 5.4 of the SI we report a steady state analysis with respect to *δ* and *α* which mirrors the analysis done for the negative feedback architecture. We find that increasing the positive feedback rate *δ* increases the percent activity of each *T_i_* in all topologies; interestingly, for the handshake/neighbor topologies the flux mismatch is worsened with a large *δ*. The response time for each *T_i_* is generally large (above 30–50 minutes), and improves for large *α* and *δ*.

This positive feedback architecture may be implemented using transcriptional circuits as done for the negative feedback system. We propose a plausible design scheme in Section 4.2 of the SI, together with numerical simulations listing all the expected reactions. While plausible, this design suffers from undesired self-inhibition pathways unavoidable with the proposed design. Preliminary experiments on this system^30^ (not reported in this manuscript) highlight the need for improved reaction mechanisms with tighter control over such undesired reactions.

## 3 Conclusions and Discussion

We have described the use of negative feedback as a mechanism to match production and demand in biochemical networks, and we provided an experimental demonstration of its effectiveness using synthetic transcriptional system *in vitro*.^19, 22, 23^ We identified “demand” as a target ligand or binding site that sequesters the output of a molecular device: in the context of our implementation, we considered artificial “genelets” whose RNA outputs bind to downstream target RNA species. In the absence of regulation, uncertainty in the demand or in the production rate of the molecular device output can cause imbalances between the concentration of available and consumed output. This imbalance can in turn result in accumulation of undesired reactants in a network, and result in malfunction of a device otherwise performing well in isolation. We show that negative autoregulation provides several advantages, in particular minimization of unused output of a device and robustness of its activity level relative to uncertainty in the output production rate. We also find that negative feedback helps reducing the sensitivity of the available output fraction with respect to uncertain downstream “load” (demand) concentration: these results are consistent with the role of negative feedback in retroactivity theory.^33^ However, unlike the typical retroactivity theory setting, we consider a “consumptive” load binding mechanism (*i. e.* the load binds irreversibly to the output), and we do not include an output amplification “gain” as part of our feedback scheme.

The ability of negative feedback to automatically tune activity as a function of downstream demand is particularly relevant when the outputs of multiple devices interact to create possibly complex functionalities or assemblies. Uncertainty and variability of molecular demand would be significant challenges that careful open-loop tuning of each device would not address. We considered a minimal, two-elements network where the outputs interact to form a product, and excess of either output is designed to downregulate its own production. We designed a transcriptional network where the RNA transcripts of two synthetic genes are complementary and bind to form an inert product; however, excess of either RNA species self-inhibits by promoter displacement. Our assays show that, as expected, negative feedback balances production and demand in the synthetic genes, leveling their activity to comparable levels. Finally, through numerical analysis we examined the scalability of our system to networks of *n* devices, identifying three possible topologies of output interconnection. Negative autoregulation still guarantees a matched flux of outputs for all topologies, although topologies with a larger number of interconnections achieve faster response times, and stationary activity and relative flux mismatch are more easily tunable for each device as a function of the negative feedback reaction rates.

Through numerical simulations we contrasted negative autoregulation with a cross-activation scheme. Our analysis suggests that this positive feedback scheme is effective in matching and maximizing production rates within a network, and it would be thus appropriate for products in high demand.^10^ However, its experimental implementation using transcriptional networks is challenging (as discussed in Section 4.2 of the SI) due to the presence of undesired self-inhibitory interactions not easily avoidable by design; these unwanted reactions may be eliminated using “translator” DNA gates.^41, 42^ Again through simulations, we showed that our cross-activation scheme can achieve matched production and demand in larger networks, but the number of required regulatory pathways scales poorly with the number of devices. In addition, our analysis for networks with 2 and 3 interconnected devices highlights that positive feedback slows down the network response time (relative to a negative autoregulation-based network with consistent parameters). This observation agrees with the slow response time introduced by positive feedback in transcriptional control of gene expression,^43^ and on the delay-inducing behavior of feedforward loops.^43^

Our experimental implementation using transcriptional circuits shows the viability of the negative autoregulation scheme in the context of *in vitro* networks.^24^ Transcriptional circuits have been used as a toolbox to build a variety of devices including toggle switches,^19^ memory elements,^44^ oscillators,^22, 23^ and a variety of other network motifs.^45, 46^ These circuits are easily programmable and expandible: regulatory interactions are designed through nucleic acid strand displacement and hybridization cascades, whose thermodynamics and kinetics can be predictably tuned by optimizing their base pair content^47^ with a variety of software toolboxes.^25, 26^ Rationally programmed nucleic acid networks can be easily interfaced with an array of ligands and physical signals through aptamers.^48, 49^ Thus, the significance of our experimental implementation goes beyond the proof of a principle: systematic use of negative autoregulation in the context of complex synthetic *in vitro* DNA networks will improve their robustness and adaptability to uncertainty in the environment. In particular, our scheme may be immediately used in the context of regulated, cotranscriptional production of RNA self-assembled structures,^5, 6^ where mismatched production and demand of components can favor the formation of incorrect complexes.

The bottom-up construction of dynamic molecular devices is a tremendous opportunity to both improve our understanding of natural biological functions and create new, artificial biotechnologies. Negative feedback has been widely used to design and tune the dynamics of synthetic *in vitro* devices such as oscillators and bistable systems.^19, 22, 23, 50^ We envision that negative feedback will also be needed to guarantee functionality when multiple devices are integrated in large scale networks, possibly requiring hyerarchical, layered feedback loops akin to modern networked control systems.^7^ Negative autoregulation mechanisms similar to the architecture described in this work will be useful not only to automatically match production and demand of individual biochemical production processes, but also to guarantee modular and adaptive input-output behaviors of components within a complex interconnected system.

## 4 Materials and Methods

### 4.1 DNA oligonucleotides and enzymes

All the strands were purchased from Integrated DNA Technologies, Coralville, IA. Genelets were labeled with TAMRA and Texas Red at the 5′ and of their non-template strands; activators were labeled with the IOWA black RQ quencher at the 3′ end. For transcription experiments we used the T7 Megashortscript kit (#1354), Ambion, Austin, TX which includes a proprietary T7 RNA polymerase enzyme mix. *E. coli* RNase H was purchased from Ambion (#2292).

### 4.2 Oligonucleotide sequences

Sequences are reported in section 1.2 of the SI file.

### 4.3 Transcription

Genelet templates were annealed with 10% (v/v) 10*×* transcription buffer-part of the T7 Megashortscript kit (#1354) - from 90°C to 37°C for 1 h 30 min at a concentration 5–10*×* the target concentration. The DNA activators were added to the annealed templates from a higher concentration stock, in a solution with 10% (v/v), 10*×* transcription buffer, 7.5 mM each NTP, 4% (v/v) T7 RNA polymerase, and .44% (v/v) *E. coli* RNase H. Each transcription experiment for fluorescence spectroscopy was prepared for a total target volume of 70 *µ*l. Samples for gel studies were quenched using a denaturing dye (80% formamide, 10 mM EDTA, 0.01 g XCFF).

### 4.4 Data acquisition

The fluorescence was measured at 37°C every two minutes with a Horiba/Jobin Yvon Fluorolog 3 system. Excitation and emission maxima for TAMRA were set to 559 nm and 583 nm, respectively, according to the IDT recommendation; for Texas Red the maxima for the spectrum were set to 598–617 nm. Raw fluorescence data Φ(t) were converted to estimated switch activity by normalizing with respect to maximum fluorescence Φ_max_ (measured before adding activators and enzymes) and to minimum fluorescence Φ_min_ (measured after adding activators and before adding enzymes):

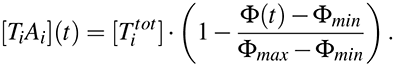

For the adaptation experiments, normalization was done by measuring maximum and minimum fluorescence levels at the beginning of the experiment, and assuming that the maximum fluorescence level scales linearly with the change in total fluorescently labeled strands, while the minimum is not significantly affected by that variation. We used the formula:

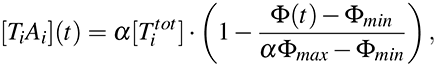

where *α* is a factor that scales the total amount of template as it varies in the experiment.

Denaturing polyacrylamide gels (8% 19:1 acrylamide:bis and 7 M urea in TBE buffer, 100 mM Tris, 90 mM boric acid, 1 mM EDTA) were run at 67°C for 45 min with 10 V/cm in TBE buffer. Samples were loaded using Xylene Cyanol FF dye. For quantitation, denaturing gels were stained with SYBR Gold (Molecular Probes, Eugene, OR; #S-11494). As a reference, we used a 10-base DNA ladder (Invitrogen, Carlsbad, CA; #1082-015). Gels were scanned using the Molecular Imager FX (Biorad, Hercules, CA) and analyzed using the Quantity One software (Biorad, Hercules, CA).

### 4.5 Numerical simulations

Numerical simulations were run using MATLAB (The MathWorks). Ordinary differential equations were integrated using the `ode23` routine. Data fitting was performed using the `fmincon` routine. Details on the data fitting procedure are in Section 1.6.4 of the SI.

## Acknowledgement

The authors would like to thank Jongmin Kim, Paul Rothemund, Franco Blanchini, and Erik Winfree for their feedback. In particular we thank Erik Winfree for sharing laboratory facilities. This work has been supported by the National Science Foundation through grants CCF-0832824 (“The Molecular Programming Project”) and CMMI-1266402; and by the Institute for Collaborative Biotechnologies through grant W911NF-09-0001 from the U.S. Army Research Office.

